# Real-time analysis of the diphtheria outbreak in forcibly displaced Myanmar nationals in Bangladesh

**DOI:** 10.1101/388645

**Authors:** Flavio Finger, Sebastian Funk, Kate White, Ruby Siddiqui, W John Edmunds, Adam J Kucharski

## Abstract

**Background:** Between August and December 2017, more than 625,000 Rohingya from Myanmar fled into Bangladesh, settling in informal makeshift camps in Cox’s Bazar district, joining 212,000 Rohingya already present. In early November, a diphtheria outbreak was reported in the camps, with 440 cases being reported during the first month. A rise in cases during early December led to a collaboration between teams from Médecins sans Frontières – who were running a provisional diphtheria treatment centre – and the London School of Hygiene & Tropical Medicine with the goal to use transmission dynamic models to forecast the potential scale of the outbreak and the resulting resource needs.

**Methods:** We first adjusted for delays between symptoms onset and case presentation using the observed distribution of reporting delays from previously reported cases. We then fit a compartmental transmission model to the adjusted incidence stratified by age-group and location. Model forecasts with a lead-time of two weeks were issued on 12th, 20th, 26th and 30th December and communicated to decision-makers.

**Results:** The first forecast estimated that the outbreak would peak on 16th December in Balukhali camp with 222 (95% prediction interval 126–409) cases and would continue to grow in Kutupalong camp, requiring a bed capacity of 200 (95% PI 142–301). On 16th December, a total of 70 cases were reported, lower than forecasted. Subsequent forecasts were more accurate: on 20th December we predicted a total of 701 cases (95% PI 477–901) and 105 (95% PI 72–135) hospitalizations until the end of the year, with 616 cases actually reported during this period.

**Conclusions:** Real-time modelling enabled feedback of key information about the potential scale of the epidemic, resource needs, and mechanisms of transmission to decision-makers at a time when this information was largely unknown. By December 20th, the model generated reliable forecasts and helped support decision-making on operational aspects of the outbreak response, such as hospital bed and staff needs, and with advocacy for control measures. Although modelling is only one component of the evidence base for decision-making in outbreak situations, suitable analysis and forecasting techniques can be used to gain insights into an ongoing outbreak.

## Background

Between August and December 2017, more than 625,000 Rohingya fled into Bangladesh as a result of large scale operations conducted by the Myanmar military in Rakhine state. This resulted in one of the largest refugee crises in recent history. The new refugees joined more than 212,000 Rohingya already present from past exoduses and settled in mostly informal makeshift camps and amongst the host community [1]. The poor living conditions typically seen in refugee settings – such as reduced access to healthcare, low standards of water, sanitation and hygiene (WASH), malnutrition and high population density – are often associated with infectious disease outbreaks. Such settings can enable transmission of infections associated with poor water and sanitation, such as cholera and hepatitis E [2, 3], as well as infections that in other settings are prevented through routine childhood vaccination, such as measles and diphtheria [4].

On 10th November, a case of diphtheria was reported to a health care facility in Balukhali run by Médecins sans Frontières (MSF). Diphtheria is caused by the diphtheria toxin producing bacterium Corynebacterium diphtheriae, which is transmitted through droplets and close physical contact, typically resulting in disease of the upper respiratory tract. Symptoms can include the formation of a pseudo-membrane obstructing airways or markedly enlarged lymph nodes. Common complications include difficulty breathing and swallowing and myocarditis. The incubation period is typically between 2–5 days (range 1-10) [5], with an estimated basic reproduction number of 4-5 [6]. Due to its high transmissibility and reported case-fatality rates of over 10% [5], diphtheria was a worldwide major public health concern with 1 million cases and 50,000 to 60,000 deaths per year in the 1970s, leading to the inclusion of diphtheria toxoid-containing vaccines in the Expanded Programme on Immunization (EPI) by the World Health Organization (WHO). As a result, the global diphtheria incidence has decreased drastically in the second half of the last century (by over 90% between 1980 and 2000), but remains of significant concern in areas with low vaccination coverage [5]. Recently, outbreaks have occured in Yemen, Venezuela, Indonesia and Haiti [7, 8].

In the month after the first case was reported in Balukhali, there were 440 additional suspected cases reported in nearby refugee settlements, 168 of which were reported on 9th Dec 2017 alone. An initial temporary Diphtheria Treatment Centre (DTC) in Balukhali run by MSF opened in the week starting on the 17th December (epidemic week 51). In the early stages of an infectious disease outbreak, it is crucial to understand the epidemiology of the infection. By quantifying transmission dynamics, it is possible to produce forecasts of future incidence [9, 10] and evaluate the potential impact of control measures [11, 12].

The significant rise in the number of diptheria cases in early December (Figure 1) led to the establishment of a collaboration between teams from MSF and the London School of Hygiene & Tropical Medicine (LSHTM) with the goal to forecast the potential scale of the outbreak and the resulting resource needs using transmission dynamic models. The first forecast was issued on 12th December, with three more subsequently issued, before the DTC in Balukhali closed on 8th January 2018 (handing over diphtheria activities to several newly-opened DTCs run by different organizations). Such analysis can face multiple challenges in real-time, including delays and variability in available data streams, limited pre-existing epidemiology studies, and knowledge gaps about risk factors and immunity in the host population. As well as describing the modelling methodology and forecasts, we report on the practical implications of the analysis, examining the role real-time modelling of infectious disease dynamics can play in operations and decision-making in a complex humanitarian crisis.

**Figure 1:**
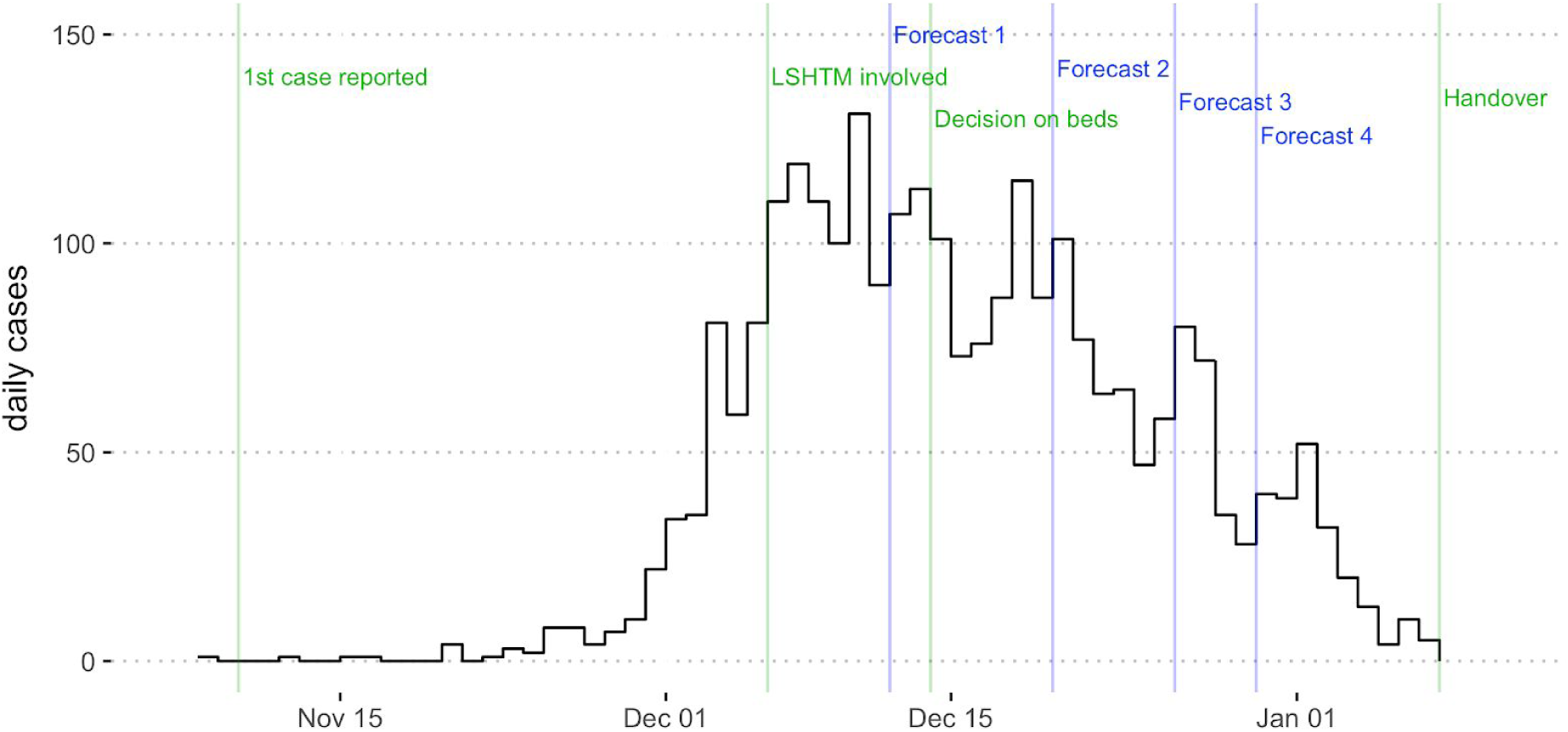
Outbreak analysis timeline with respect to the epidemic curve (black line). Green lines show timing of events relevant to analysis: reporting of first case; involvement of modellers at LSHTM; MSF decision on bed numbers required; MSF handover of treatment centre. Blue lines show date on which each of the four LSHTM forecasts were communicated to MSF.

## Methods

### Data

Between 8th November and 31st December 2017, a total of 2,624 cases (495 from Kutapalong, 1,868 from Balukhali and 261 from other or unknown nearby locations) presented at the Diphtheria Treatment Centre in Balukhali run by MSF. The total refugee population had been estimated at around 608,000 in early December (Additional File 1). From 9th December 2017 to 12th January 2018, we received daily line lists of suspected cases seen at this centre (Additional File 2). Fields included patient identification number, sex, age, approximate address of patient, date of onset of symptoms, date of reporting to the DTC, signs and symptoms, treatment and clinical outcome. First, we checked the line list for objectively erroneous values – such as dates that were in the future or dates of reporting prior to date of onset of symptoms – and corrected these where possible. We then computed the daily crude incidence within three age groups (0-4, 5-14 and >=15 years) and two geographical locations (Balukhali and Kutupalong, with other/unknown locations omitted from the analysis).

### Adjustment for delayed reporting

To adjust for delays between symptoms onset and case presentation, and estimate the actual incidence at a given point in time, we computed the cumulative distribution of reporting delays of reported cases (defined as the number of days between symptom onset and case presentation). We then divided these crude daily incidence values by the corresponding values in this distribution (e.g. delay 0 for the current day, delay 1 for the previous day etc.) to obtain the adjusted incidence. Initially we used all previously reported cases to compute this delay distribution, but changed the analysis window as more data became available. From 18th December onwards we used cases with symptom onset between 10th December and 16th December to compute the delay distribution; from 24th December we used cases with symptom onset between 17th December and 23th December, and starting on 30th December, we used all cases with symptom onset since 10th December.

### Mathematical model and forecasting

To model the epidemic, we used a deterministic Susceptible-Exposed-Infected-Recovered (SEIR) transmission model (Additional File 3), which allowed for different transmission rates in each location and different levels of susceptibility in each age group. An additional compartment represented the number of patients currently hospitalised as a result of diphtheria infection.

The gaps in time between the onset dates of early reported cases suggested the generation time of infection may have been around 4–5 days (Additional File 3). We therefore assumed an incubation period of 3 days and an infectious period of 3 days; this implied an expected generation time of 4.5 days. Vaccination coverage for DPT3 in Myanmar was reported to be 85% in 2012 [13], but coverage was likely to be lower in the Rohingya population [14]. In November 2017, a health survey performed by MSF estimated that measles vaccination coverage in Rohingya children aged between 6 to 59 months was 20–25%, following a vaccination campaign in children between 9 months and 14 years old [15]. Vaccination data for diphtheria were not available, but we assumed that 20% of the 5–14 age group were initially immune to diphtheria.

As well as the fixed parameters, the model had seven parameters that were estimated independently for each location: the proportion of cases reported, the transmission rate, the relative susceptibility in under 5 and over 14 age groups, and the initial proportion of infectious in each age group. When an outbreak is growing exponentially, it is not possible to jointly estimate the initial number of infectious cases and the proportion of cases reported because the two parameters are inversely correlated. To obtain a prior distribution for the proportion of cases that might be reported, we performed a rough calculation using data from the pre-vaccination era. Prior to widespread DTP3 vaccination coverage in the UK, there were around 55,000 cases of diptheria per year [16] and 750,000 live births each year [17]. Given diphtheria has a relatively high basic reproduction number, R_0_, of 4-5, almost all initially susceptible individuals would be expected to eventually become infected in the absence of vaccination [6]. Hence if there were 750,000 live births per year, almost all of these would become infected at some point. Based on the data from the pre-vaccination era, with 55,000 annual cases reported, this would suggest that at least 7% (55,000/750,000) of diphtheria infections appear as cases. We therefore imposed a strong gamma prior distribution on the proportion of cases reported, which had a mean of 10% and a standard deviation of 2.2%. We assumed log-uniform priors on all other parameters. The fourteen free parameters in the model (seven for each camp) were calibrated to the adjusted incidence at each location using a Markov Chain Monte Carlo (MCMC) model fitting procedure. A summary of model parameters is given in Table 1.

**Table 1:** Parameters used in the model. Parameters that are camp-specific take independent values for Balukhali and Kutupalong, and the initial proportion of people infectious is specific to each age group in each camp.

| Parameter | Age or camp specific? | Value | Prior distribution |
| --- | --- | --- | --- |
| Incubation period | No | 3 days |  |
| Infectious period | No | 3 days |  |
| Transmission rate | Camp | Fitted | Log-uniform(0,∞) |
| Proportion of cases reported | Camp | Fitted | Gamma(20,0.005) |
| Dispersion parameter for reporting | Camp | Fitted | Log-uniform(0,∞) |
| Relative susceptibility in under 5 age group | Camp | Fitted | Log-uniform(0,1) |
| Relative susceptibility in over 15 age group | Camp | Fitted | Log-uniform(0,1) |
| Initial proportion susceptible | No | 0.85 |  |
| Initial proportion infectious | Camp and age | Fitted | Log-uniform(0,1) |

The model was used to generate forecasts of future incidence on December 12th, December 20th, December 26th and December 30th. To produce a forecast, the model was calibrated to the past adjusted incidence in each age group and location and 1000 epidemics simulated up to two weeks into the future. Median and uncertainty (95% prediction intervals (PI)) were communicated to partners. When forecasting bed requirements, we assumed that 15% of reported cases would require treatment as inpatients, and an average hospital stay would be 5 days. These estimates were informed by early patient data in the line list.

## Results

### Adjustment for delayed reporting

The delay between symptom onset and case presentation was 2 and 6 days respectively for the first two reported cases. This subsequently increased to 13 (range 5 to 21) days before stabilizing around a median value of 2 (range 0 to 12) days from early December onwards (Figure 2A). Comparing estimated incidence adjusted for reporting delays with the final incidence subsequently reported, we found that our method reduced the bias introduced by delayed reporting, although results showed high variability (Figures 2B and C).

**Figure 2:**
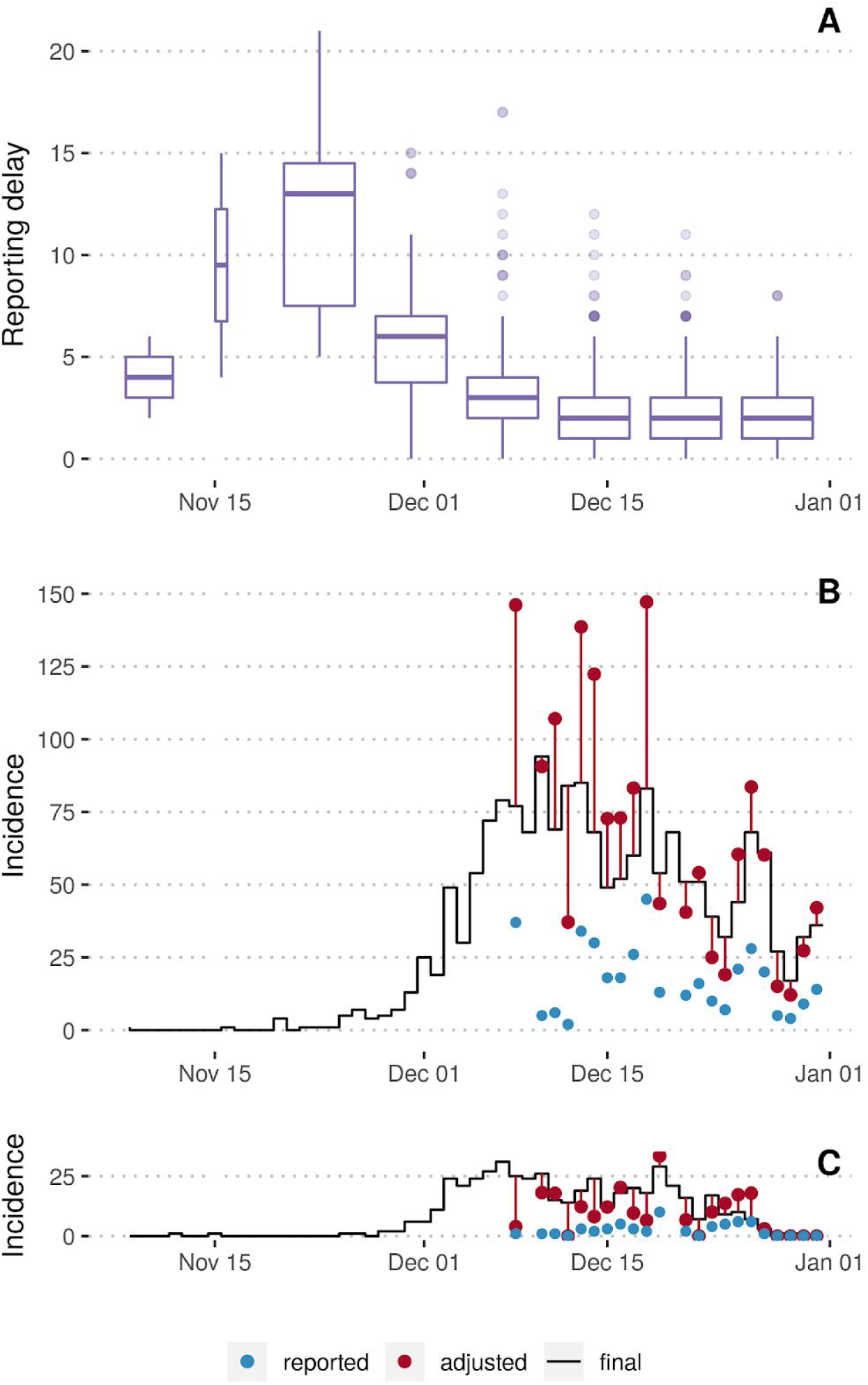
Adjustment for delay between symptoms onset and case presentation (reporting delay). Evolution of the reporting delay (vertical axis) by epidemic week (horizontal axis) (A). Daily incidence of diphtheria cases in Balukhali (B) and Kutupalong (C) as reported within the first day after symptoms onset (blue dots), adjusted for reporting delays (red dots) and as seen retrospectively (black line, data from 12 January 2018).

Between 8th November and 31st December, a median of 31% (95% PI 4–48%) of cases were reported within one day of symptoms onset (i.e. until the end of the first day after onset), 61% (95% PI: 22–82%) within two days and 84% (95% PI: 45–94%) within three days.

Adjusting for delayed reporting we overestimated daily incidence by a median of 7% (95% PI: −61 to 46%) from the unadjusted incidence known within one day after symptom onset, by 5% (95% PI: −27 to 52%) from the unadjusted incidence two days after onset and by 0% (95% PI: −17 to 40%) from the unadjusted incidence three days after onset. Note that negative percentages indicate underestimation of the actual incidence here.

### Forecasts

We generated four sequential forecasts of future incidence, each stratified by age group and location. The initial forecast was produced and communicated to MSF field staff on 12th December. Subsequent forecasts were issued on December 20th, 26th and 30th (Figure 3). In the first forecast, we estimated that the epidemic would peak on 16th December in Balukhali with 222 (95% PI: 126–409) cases and would continue to grow beyond the 2 weeks forecast horizon in Kutupalong (Figure 4). The required bed capacity at the end of the forecast horizon was estimated to be 200 (95% PI: 142–301). In reality, the peak of the epidemic (131 reported cases, 94 of which in Balukhali) had already occurred on 10th December, although it was not possible to conclude this from the data available in real-time. On 16th December, a total of 70 cases were to be reported, lower than forecasted.

**Figure 3:**
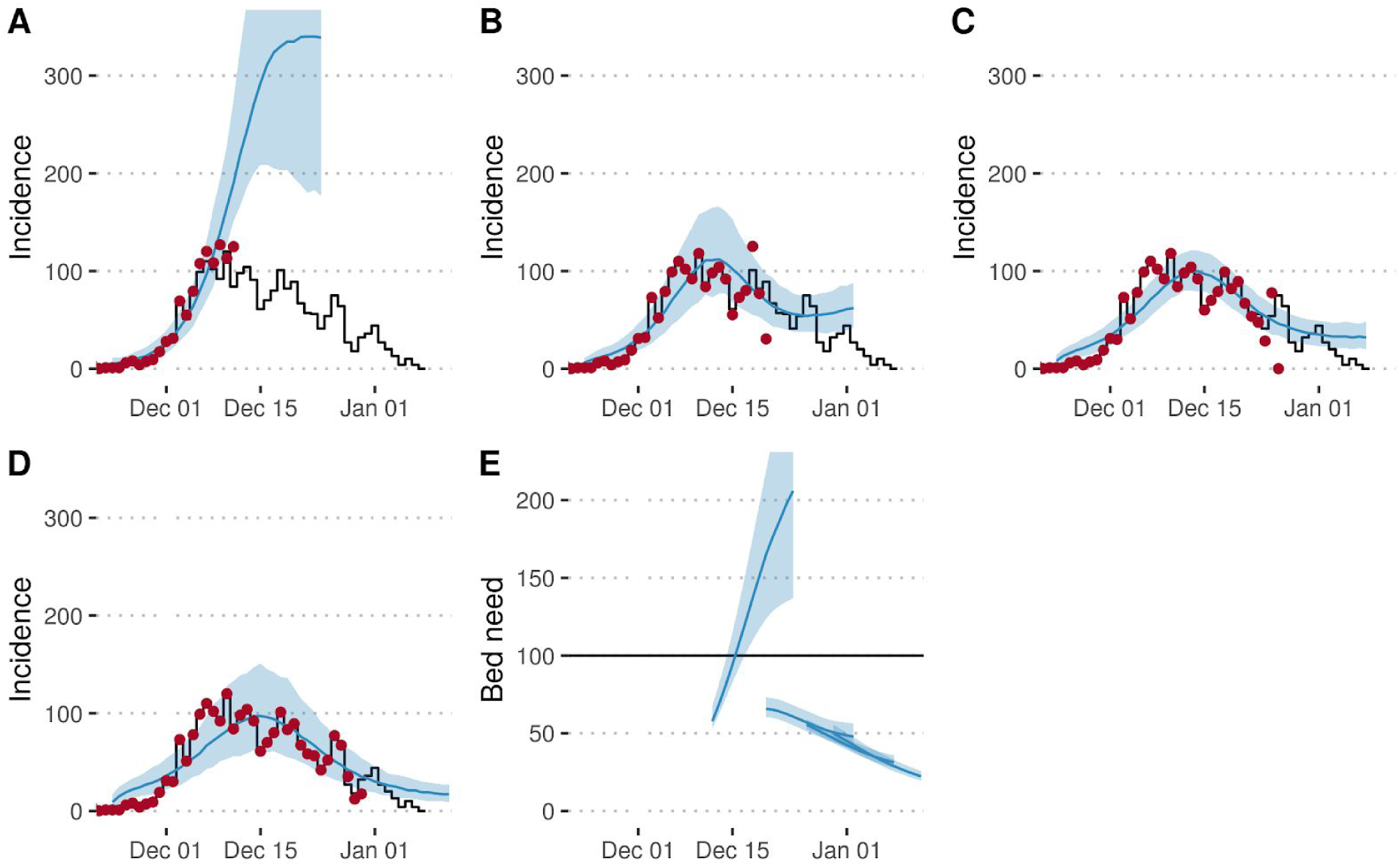
Total incidence over all age groups and locations (A-D) and bed need as forecasted by the model. Black lines show data as reported by 12 January 2018, red dots the adjusted incidence and blue lines and shaded areas the median and 2.5% and 97.5% percentiles according to 1000 model runs forecasting from 12th December (A), 20th December (B), 26th December (C) and 30th December (D). Forecasts of bed need issued on the same dates (E). The horizontal line shows the number of beds provided as of a decision taken on 14th December.

**Figure 4:**
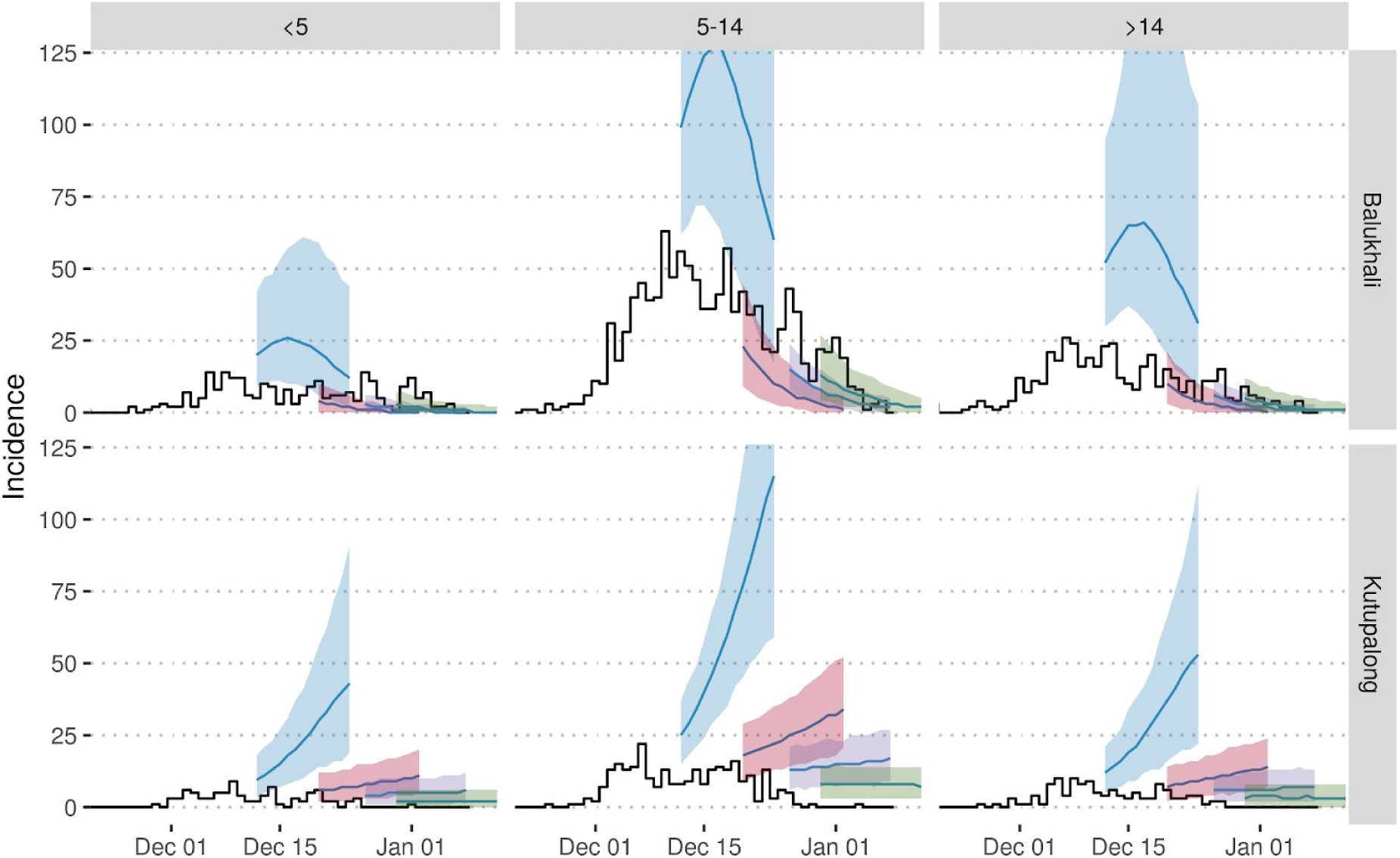
Incidence by location (rows) and age group (columns) as forecasted by the model. Black lines show data as reported by 12 January 2018 and coloured lines and shaded areas the median and 2.5% and 97.5% percentiles according to 1000 model runs forecasting from 12th December (blue), 20th December (red), 26th December (purple) and 30th December (green).

Forecasts became more accurate later during the epidemic. On 20th December, the model predicted a total of 701 cases (95% PI 477–901, this corresponds to 105 hospitalized patients under our assumptions) and on 26th December a total of 245 cases (95% PI 169–329, this corresponds to 37 hospitalized cases) up to the end of the year. In reality, 616 and 252 cases were actually reported during those respective periods. According to our model, R_0_ was equal to 8.2 (95% PI 7.2–9.8) in Balukhali and 5.8 (95% PI 5.0–6.9) in Kutupalong based on the initial forecast. Estimates later stabilized at lower values of 7.6 (95% PI 7.1–8.1) and 3.3 (95% PI 3.0–3.6) respectively on 26th December. The proportion of cases reported was estimated to be significantly below the assumed prior median value of 10% (Figure 5). In both camps, susceptibility in the under 5 and over 15 age groups was estimated to be at least 50% lower than susceptibility in the 5–14 age group (Figure 5).

**Figure 5:**
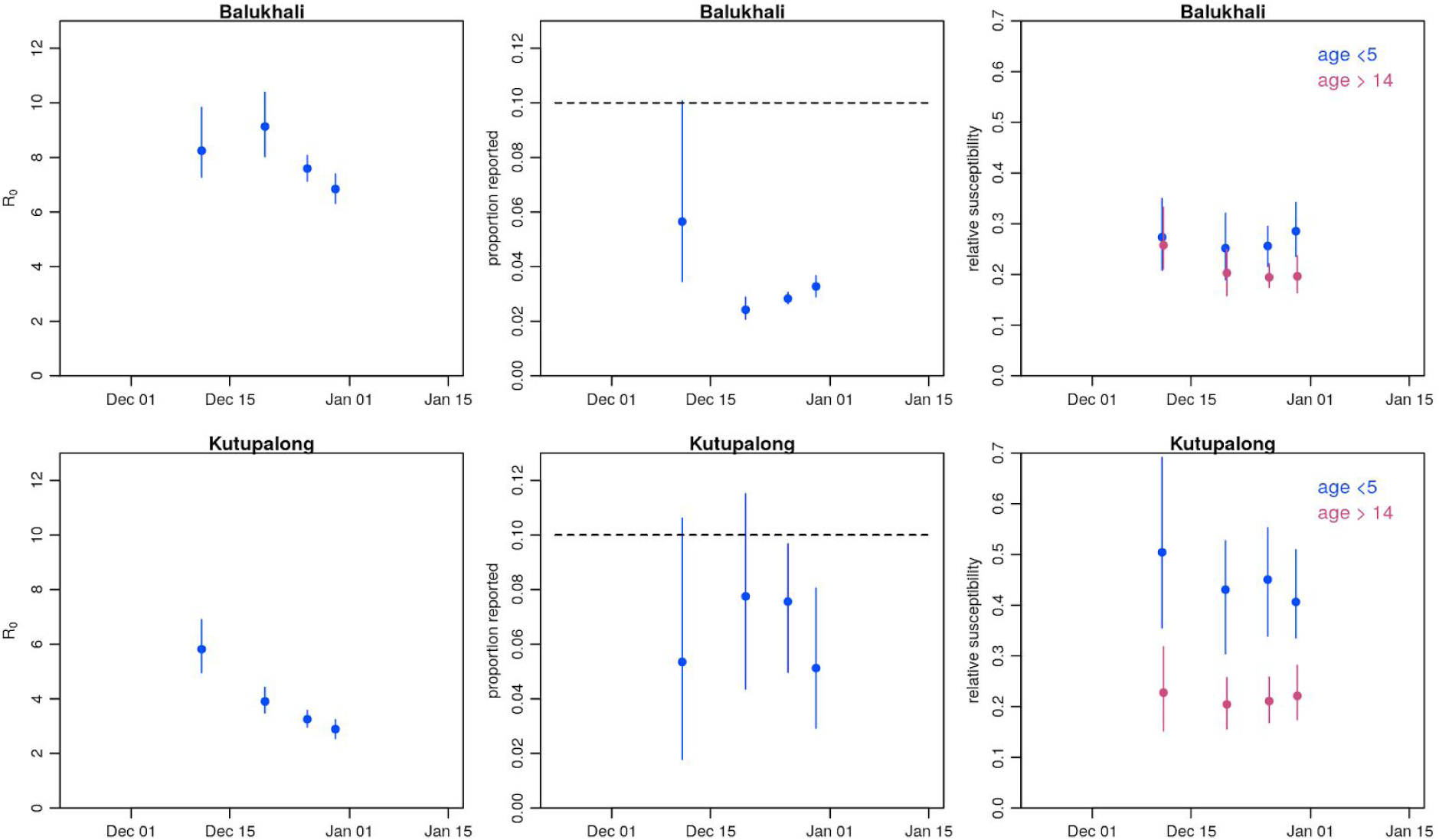
Posterior parameter values. Posterior ranges (vertical lines) and median values taken by model parameters for forecasts done on 12th December, 20th December, 26th December, 30th December and 8th January. The horizontal dashed lines show the mean value of the prior used for the proportion of reported. Uniform priors were used for other parameters.

### Operations and decision making

The forecasts contributed to an evidence base that helped support operational aspects of the response, as well as advocacy for control measures. During December, staffing was increased in response to the outbreak. MSF employed a strategy of surge staffing for international staff and expedited recruitment of national staff doctors and nurses. On 17th December a conservative decision to make a total of 100 hospital beds available was taken by MSF, using the high number of potential cases forecasted by the real-time modelling to guide the decision (with a view to monitor the modelling outputs over the coming weeks). It was also decided to categorise beds into two severity levels depending on clinical signs [18] and to treat mild cases in the community, which helped ensure that the available number of beds was never exceeded. Efforts to trace contacts of patients were intensified. Stocks of diphtheria antitoxin, which was in a global shortage due to other outbreaks in Yemen, Venezuela, Indonesia and Haiti at the time, antibiotics and other supplies were increased.

Initial advocacy for vaccination had centered on a broad age group. The modelling analysis highlighted that the under 5 and over 14 age groups were less susceptible relative to the 5–14 age group. As a result, the 5-14 year-old group contributed most to disease transmission, likely as a result of lack of vaccination in the displaced population before their arrival in Bangladesh. The discussions around forecasting also contributed to advocacy to scale up outbreak response by other actors such as the Global Outbreak Alert and Response Network (GOARN), Samaritan's Purse and the UK’s Emergency Medical Team, and helped lead to a closer collaboration between key partners such as MSF and WHO.

## Discussion

In this study, we have shown how transmission dynamic models and forecasting techniques provided insights into the epidemiological processes underlying the diphtheria outbreak in Forcibly Displaced Myanmar Nationals living in camps and makeshift settlements in Cox’s Bazar district, Bangladesh. This enabled real-time analysis to estimate the course of the outbreak and corresponding resource needs.

Although our model captured the overall dynamic of the epidemic, there were several limitations to the modelling analysis. A number of key epidemiological parameters were unknown and had to be assumed from the literature or inferred from incidence data. In addition, the adjustment for reporting delays was initially biased upwards as delays between onset and case presentation shortened significantly during the early epidemic. These factors, combined with parameter uncertainty and a rapid increase in cases during this time, led to the first forecast overestimating the number of future cases, and made it difficult to capture the dynamics in Kutupalong.

Another limitation in the early stages of the analysis was that some key epidemiological unknowns could not be estimated. During the initial exponential growth phase of an epidemic it is not possible to jointly estimate all key unknown transmission and reporting parameters. As a result, it is necessary to impose prior assumptions on at least some of these parameters; we constrained prior susceptibility and reporting rate. Such assumptions, combined with remaining uncertainty about unknown parameters, can lead to substantial variability in forecast trajectories and potential bias in model outputs. In addition, the deterministic model we used did not capture time-variations in key parameters such as the reproduction number (i.e. due to interventions, such as contact tracing and active case finding or the WHO-lead vaccination campaign initiated on 12th December 2017) or the reporting rate (i.e. due to changes in health-seeking behaviour induced by health promotion activities and circulating information about the outbreak itself). The model attributed any uncertainty to the fitted parameters and the reporting process, rather than stochasticity in transmission. A stochastic model could have been used to include a more accurate representation of uncertainty and to capture time variations in parameters. However, such a model would have been more time consuming to set-up and calibrate, and would still have been reliant on imperfect data.Like the deterministic model, it would also have required strong assumptions to be made about several aspects of the epidemiology. Moreover, the introduction of vaccination is unlikely to have had an impact within the time frame analysed here given the delay to protection, incubation period following infection, and the delay in reporting following onset. Finally, we assumed a fixed population size, though in reality there can be a substantial influx of people into camps during outbreaks, as well as movement within camps; understanding how such movements might affect outbreak dynamics in general would be worth investigating in future studies.

Our estimated values for the basic reproduction number R_0_ were in agreement with values from the literature [6, 19] and other estimates for the same epidemic [20], although our assumed generation times were lower and estimates of the reporting rate were higher compared to an analysis of the early diphtheria outbreak dynamics by Matsuyama et al. [20], which did not stratify by age or camp. Whereas we assumed that the susceptibility was greatest in the age group of 5-14 years, the proportion of cases in the age group of 15 years and above was higher before the epidemic peak than after. This may indicate either that adults made a substantial contribution to transmission during the epidemic growth phase [21], or that relative age-specific reporting changed during the course of the outbreak.

Construction of mechanistic epidemic models makes it possible to formalize assumptions about the epidemiological processes underlying an outbreak, incorporating expert knowledge and context-specific analysis of the local situation. When working in real-time, the main challenge lies in quickly consolidating all necessary information – in an often complex and variable emergency situation – to be able to make appropriate assumptions in a model. In our case, a better understanding of epidemiological processes, disease characteristics, case reporting, and prior vaccination status would have allowed for more accurate assumptions and potentially more accurate forecasts. Despite regular discussions between LSHTM and MSF during December, the first forecasts were only delivered a month after the first case was reported.

Our experiences of real-time modelling and analysis during this outbreak highlighted the importance of effective ongoing communication with field staff. As well as enabling access to real-time data (including incidence, demography and geography), staff can also provide additional context and information such as: the general epidemiological situation, likely vaccination status of the population, the nature and severity of symptoms, health seeking behaviour and access to health care, the sanitary situation, and population movements. To maximise the future benefit of real-time modelling, it would be advantageous to build strong, long-term collaborations between organizations providing outbreak responses and modellers [22]. Such collaborations should focus on establishing well-defined processes (i.e. analysis pipelines) on how to collect, treat and share relevant data and other information from the field with modellers, ideally embedding an experienced modeller or data manager in the outbreak response team, and enabling model results and model-based recommendations to be fed back to field staff and decision-makers, whose input can in turn inform subsequent analysis.

## Conclusions

Although modelling is only one component of the evidence base for decision-making in outbreak situations, we have shown that suitable analysis and forecasting techniques can be used to gain insights into an ongoing outbreak.

In the context of the diphtheria outbreak in Bangladesh, real-time modelling made it possible to feedback key information about the potential scale of the epidemic, likely resource needs and underlying mechanisms of transmission to decision makers at a time when this information was largely unknown. By December 20th, our model was able to generate reliable forecasts with a lead-time of two weeks.

We advocate that such analysis can be further developed in the future through strengthening collaborations and setting up bi-directional data and information flow pipelines linking modellers with decision-makers and field staff, so that real-time modelling can rapid and routine contributions to outbreak response.

## Abbreviations

WASH: Water, Sanitation and Hygiene
MSF: Médecins sans Frontières
WHO: World Health Organization
LSHTM: London School of Hygiene and Tropical Medicine
GOARN: Global Outbreak Alert and Response Network
DTC: Diphtheria Treatment Centre
MCMC: Markov Chain Monte Carlo

## Declarations

### Ethics approval and consent to participate

This research fulfilled the exemption criteria set by the MSF Ethics Review Board for a posteriori analyses of routinely collected clinical data and thus did not require MSF Ethics Review Board review.

### Consent for publication

Not applicable

### Availability of data and material

All data generated or analysed during this study are included in this published article [and its supplementary files].

### Authors’ contributions

FF, AJK, SF and WJE designed the methodology, performed the analysis and interpreted the results. RS and KW acted as liaison with field staff from MSF, provided data and in-situ information, were involved in interpreting the results and incorporated model results into the operational decision making processes. FF and AJK wrote the first draft of the manuscript manuscript and all authors read and approved the final manuscript.

## Acknowledgments

We thank Crystal Van Leeuwen and John Guzek as well as all other MSF staff who participated in the data collection and outbreak response. We would also like to thank all partner organizations involved in the outbreak response, and in particular the Directorate General of Health Services of Bangladesh, for the excellent collaboration.

## Additional Files

**Additional File 1** (CSV table)

Population data

**Additional File 2** (CSV table)

Incidence data - The unadjusted incidence data extracted from linelists and used for this analysis.

**Additional File 3** (PDF)

Supplementary Information - Detailed description of the model as well as the model fitting procedure, including equations and posterior parameter estimates.

